# HiCTMap: Detection and Analysis of Chromosome Territory Structure and Position by High-throughput Imaging

**DOI:** 10.1101/185942

**Authors:** Ziad Jowhar, Prabhakar Gudla, Sigal Shachar, Darawalee Wangsa, Jill L. Russ, Gianluca Pegoraro, Thomas Ried, Armin Raznahan, Tom Misteli

## Abstract

The spatial organization of chromosomes in the nuclear space is an extensively studied field that relies on measurements of structural features and 3D positions of chromosomes with high precision and robustness. However, no tools are currently available to image and analyze chromosome territories in a high-throughput format. Here, we have developed High-throughput Chromosome Territory Mapping (HiCTMap), a method for the robust and rapid analysis of 2D and 3D chromosome territory positioning in mammalian cells. HiCTMap is a high-throughput imaging-based chromosome detection method which enables routine analysis of chromosome structure and nuclear position. Using an optimized FISH staining protocol in a 384-well plate format in conjunction with a bespoke automated image analysis workflow, HiCTMap faithfully detects chromosome territories and their position in 2D and 3D in a large population of cells per experimental condition. We apply this novel technique to visualize chromosomes 18, X, and Y in male and female primary human skin fibroblasts, and show accurate detection of the correct number of chromosomes in the respective genotypes. Given the ability to visualize and quantitatively analyze large numbers of nuclei, we use HiCTMap to measure chromosome territory area and volume with high precision and determine the radial position of chromosome territories using either centroid or equidistant-shell analysis. The HiCTMap protocol is also compatible with RNA FISH as demonstrated by simultaneous labeling of X chromosomes and *Xist* RNA in female cells. We suggest HiCTMap will be a useful tool for routine precision mapping of chromosome territories in a wide range of cell types and tissues.

## 1. Introduction

The organization of the eukaryotic genome extends from DNA that is coiled around histone octamers to form nucleosomes, which are further folded into a chromatin fiber, to ultimately form chromosomes [1-3]. Chromosomes are the largest unit of genome organization and are confined to distinct regions within the nucleus known as chromosome territories (CTs) [4]. The spatial organization of CTs is non-random, and CTs occupy preferred nuclear positions [5]. Changes in genome spatial organization are observed during stages of differentiation, transcription, and diseases [6-9]. Furthermore, chromosome position is cell type- and tissue-specific [10-13].

Fluorescence in situ hybridization (FISH) of entire chromosomes using specific DNA probes, also known as chromosome paints, provides vital insight into defining chromosome architecture and position within the 3D space of the nucleus [14]. By measuring the radial position of a chromosome relative to the center of the nucleus, or their relative position to each other, it has been recognized that CTs are non-randomly organized in the 3D space of the nucleus [15]. These experiments show that chromosomes have a characteristic distribution of positioning within the cell nucleus, however, the preferential positioning of chromosomes is not absolute and varies probabilistically between individual cells in a population [2, 5].

Because of the stochastic distribution of each chromosome in the cell population, accurate mapping of chromosome positions requires analysis of a relatively large number of cells. Due to limitations in imaging throughput of conventional microscopy methods, the number of cells commonly analyzed for chromosome paint experiments are often limited, typically on the order of a hundred cells or less.

We have developed HiCTMap, an unbiased and systematic method for the quantitative detection of CTs using high-throughput imaging that enables routine analysis of hundreds of nuclei and CTs per sample in an experiment. The HiCTMap protocol consists of fixation of interphase cells in 384-well plates, followed by DNA FISH staining using chromosome paint probes. Large image data sets are acquired using automated 3D confocal high-throughput microscopy and analyzed with a custom high-content image analysis workflow to determine the spatial positioning and features of chromosomes in multiple imaging channels. As proof-of-principle, we applied HiCTMap to measure the size and positioning of chromosomes 18, X, and Y in normal male and female human primary skin fibroblasts. We demonstrate that chromosome X has a more peripheral position than chromosomes 18 and Y, as previously shown by others [16-19]. Moreover, we combined RNA FISH and chromosome paint in a high throughput format to differentiate active and inactive X chromosomes and demonstrate accurate detection of the reduced size and a slightly more peripheral positioning of the inactive X chromosome.

## 2. Materials and Methods

### 2.1. Cell culture

Female (XX) and male (XY) primary human skin fibroblast were grown in DMEM media (ThermoFisher Scientific, Cat # 10566-016) supplemented with 20% FBS (ThermoFisher Scientific, Cat # 10082147), 2 mM GlutaMAX™, and 1% penicillin/streptomycin (ThermoFisher Scientific, Cat # 15140122) at 37 °C and 5% CO_2_. Cells were split 1:2 every 3-4 days and kept at a low passage (P3-5). Cells were plated in PerkinElmer Cell Carrier Ultra 384-well plates (PerkinElmer, Cat # 6057300) at a density of 4,000 cells/well and grown overnight, then fixed for 10 min in 4% PFA (Electron Microscopy Sciences, Cat # 15710), washed, and stored in DPBS (Lonza, Cat # 17-512F) at 4 °C.

### 2.2. Chromosome Paint FISH in 384-well plates

Whole chromosome paint probes were generated in house to human chromosomes 18 (HSA18), X (HSA-X), and Y (HSA-Y) using Spectrum Orange (Abbott Molecular, Abbot Park, IL, USA, Cat # 02N33-050), Dy505 (Dyomics, Jena, Germany, Cat # Dy505-dUTP), and Dy651 (Dyomics, Jena, Germany, Cat # Dy651-dUTP), for labeling of HSA18, HSA-X, HSA-Y, respectively. Detailed protocols are available online at (https://ccr.cancer.gov/Genetics-Branch/thomas-ried, Resources). Probe mix for chromosome paint probes consisting each of 200 ng - 1 μg DNA, 30 μg salmon sperm DNA (Ambion, Cat # AM9680), and 1 μg human COT1 Human DNA (Sigma-Aldrich Roche, Cat # 11581074001) per well was ethanol precipitated at −80 °C for 25 minutes and re-suspended in 12 μL of hybridization buffer per well (50% dionized formamide, pH7 (Ambion, Cat # AM9342), 20% dextran sulfate (Milipore, Cat # S4030), and 2X SSC (KSD Scientific, Cat # B3821-1000).

Fixed cells were permeabilized for 20 min in 0.5% Triton X-100 (Sigma-Aldrich, Cat # T9234-500ML) in DPBS, washed in 0.05% Triton X-100 in DPBS, incubated for 10 min in 0.1 N HCl, neutralized for 5 min in 2X SSC, and equilibrated for at least 30 min in 50% formamide in 2X SSC prior to probe addition. Probes and nuclear DNA were denatured at 85 °C for 7 min and plates were immediately moved to a 37 °C humidified chamber for overnight hybridization. The next day, plates were washed 3 times for 5 min each in 50% formamide (Sigma-Aldrich, Cat # 47670-1L-F) in 2X SSC at 42 °C, 1X SSC at 60 °C, and 0.1X SSC at 60 °C.

A DesignReady *Xist* RNA probe designed by Stellaris (25μM concentration; LGC Biosearch Technologies, Petaluma, CA, Cat # VSMF-2431-5) was diluted 1:10 in TE buffer, pH 8 and diluted again 1:3 in 12 μL of RNA hybridization buffer (10% formamide, 10% dextran sulfate in 2XSSC). Plates were hybridized at 37 °C in a humidified chamber for at least 4 hours up to overnight. Plates were then washed 3 times for 5 min with 2X SSC at 37 °C, nuclear DNA was stained with 1.25μg DAPI per well for 1 min in DPBS, and imaged. For combined DNA-RNA FISH, DNA FISH was completed as described above at 37 °C overnight with corresponding DNA FISH washes, immediately followed by RNA FISH as described above for 4 hours at 37 °C with corresponding RNA FISH washes.

### 2.3. High-throughput imaging

Imaging was performed using a 40X (0.95 NA) air objective lens on a high-throughput spinning disk confocal microscope (Yokogawa CV7000). For excitation, four lasers lines (405, 488, 561, 640 nm) were used, in conjunction with a quad-band excitation dichroic mirror (405/488/561/640 nm), a fixed short pass emission dichroic mirror (568 nm), and emission band-pass filters (filters (DAPI: BP445/45, Green: BP525/50, Red: BP600/37, and FarRed: BP676/29) in an automated and sequential mode. Image acquisition was configured to capture three-dimensional Z-stacks for each channel with 8-9 Z-sections at 0.5 μm intervals without pixel binning. The X-Y pixel size was 162.5 nm and the size of each field of view was 416 x 351 μm (2560 x 2160 pixels).12-13 fields of view were imaged per well in 15 wells per karyotype, producing approximately 15,000 16-bit images in approximately 1 hour.

### 2.4.1. Image analysis tools

The image processing steps described in the subsequent section(s) were implemented into four (4) bespoke image analysis workflows in the Konstanz Information Miner (KNIME) [20] Analytics Platform (Version 3.2.1, 64-bit) using compatible KNIME Image Processing Nodes (KNIP). We chose KNIME Analytics Platform primarily because it is open-source and readily supports development of reproducible workflows by capturing data flow. In addition, KNIME-based workflows are operating system agnostic and can scale extremely well from desktops to high-performance batch clusters.

The primary workflow configures and executes the image preprocessing and nuclear segmentation components. The primary workflow provides annotation of segmented objects (nuclei) into good and bad classes, morphological feature values, and filter training of the random forest (RF) classifier. The secondary workflow implements the training from the primary workflow for segmentation and filtration of nuclei in 2D and 3D for the plate and crops CT images from corresponding channels using the binary nucleus mask. The tertiary workflow is responsible for CT detection training. This workflow provides annotation of segmented CT objects into good and bad classes, CT intensity and morphological values, as well as training of each CT channel’s RF classifier. The quaternary workflow implements training from the tertiary workflow to segment and filter CTs (*Xist*) from multiple spectral channels in 2D and 3D while extracting parameters required for studying spatial organization and architectural features of CTs (i.e. radial positioning, area, and volume).

All KNIME workflows were either executed on a workstation running Microsoft Windows 2012 Server R2 (64-bit) with 16-cores of AMD Opteron 6212 processor (2.7 GHz) and 256GB RAM or on a dedicated high-performance batch cluster compute node running RedHat Enterprise Linux 6.9 (Santiago) with 28-cores (56-processors) of Intel X2680, 196 RAM and 300 GB of Solid State Storage.

### 2.4.2. 2D Nucleus Segmentation

The nuclei from the maximum intensity projected DAPI channel were segmented using a seeded watershed algorithm [21]. The preliminary segmentation boundaries from the seeded watershed are further refined using ultrametric contour maps (UCMs) to minimize over-segmentation [22]. Briefly, UCMs achieve this by combining several types of low-level image information (e.g., gradients and intensity) to construct hierarchical representation of the image boundaries. Under this representation boundary pixels along the nucleus periphery, typically, receive higher score than other pixels associated with internal structures of the nucleus. Applying global thresholding (e.g., Otsu [23]) on UCMs should eliminate weaker (internal) boundaries, thus, minimizing over-segmentation.

### 2.4.3. Random Forest for Filtering Overlapping Nuclei

We used a binary (two classes: *good* and *bad*) supervised classification algorithm based on RF [24, 25] to filter out under/over segmented nuclei from subsequent analysis. A 14-dimensional numeric vector describing the morphology of segmented objects (e.g., circularity, solidity, area, perimeter, and major elongation) was used for discriminating *good* and *bad* segmented objects. The first interactive KNIME workflow was used to manually annotate 100 (good-65; bad-35) segmented objects drawn randomly from images from different wells of a plate that was imaged during the plate optimization process. Subsequent nodes in the KNIME workflow then autonomously train a RF classifier on this annotated data. We used a RF classifier with 1000 decision trees, where the nodes in each decision tree were split based on the information gain ratio. This resulted in a RF classifier with out-of-bag accuracy of ~92%. The trained RF classifier was subsequently used for filtering out *bad*-class of segmented objects for all other plates referenced in this work without further (re)training.

### 2.4.4. 3D Nucleus Segmentation

For segmentation of nuclei in 3D, i.e., along the Z-stack of the DAPI channel, we first extended the 2D binary mask of nuclei along the Z-direction. This extended 3D binary mask was applied to the Z-stack of DAPI channel for extracting 3D voxels of the nucleus. The intensity voxels were then thresholded using Otsu [23] followed my morphological operations to generate the 3D binary segmentation mask of the nucleus.

### 2.4.5. 2D CT Segmentation

Prior to segmenting CTs, pixel intensity values of CT channel(s) corresponding to the 2D nucleus segmentation mask were normalized to range from 0 to 1. The normalization step was added to minimize the influence of intensity fluctuations in the CT channels between wells within a plate and across plates. Post intensity normalization, CTs in each spectral channel were segmented using the undecimated wavelet-based spot detection technique [26]. We used up to four (4) wavelet scales for segmenting CTs. The per-scale threshold factors for the algorithm were manually fixed so that CTs in all three channels (18, X, and Y/*Xist*) were reliably detected with the same set of values. These per-scale threshold factors were intentionally set low to enable detection of all CTs. Such low values, as expected, resulted in segmenting background (non-CT) regions.

### 2.4.6. Random Forest for Filtering CTs

To filter out background regions segmented by the undecimated wavelet-based spot detection technique, we again used a binary, supervised RF classifier similar to the one described earlier for filtering over/under segmented nuclei. We used an appropriate interactive KNIME workflow to randomly sample 2%-4% of the detected nuclei for each karyotype in the plate. CTs in each spectral channel were detected as described above. Our rational was to determine if the RF classifier, trained on annotated data from just one replicate, can robustly filter segmented CT objects from two biological replicates in the sample plate. If a plate contained wells with replicates, we ensured that stratified random sampling of nuclei was restricted to just one replicate. A user manually annotated correctly (class-*good CT*) and incorrectly (class-*bad CT*) segmented CT regions via the interactive nodes in this workflow – approximately 100 objects per class for each spectral channel (green, red and far-red). Next, we extracted morphometric and (normalized) intensity-based features for the objects and trained a RF classifier with 1000 decision trees and node splitting governed by information gain ratio. The out-of-bag accuracy of the RF classifiers were approximately 95.6%, 99.1% and 95.4% for CT in green, red and far-red channels, respectively. Various efforts (e.g., standardization and normalization of features) to combine three separate RF classifiers into one, generally, resulted in lower out-of-bag accuracy (data not shown). Similarly, classifier(s) trained on one plate, typically, resulted in poor filtering of incorrectly segmented (class-*bad CT*) CT regions when applied to plates with different karyotype(s). Hence, we decided to generate a separate classifier for each channel for each plate.

### 2.4.7. 3D CT Segmentation

The CTs were segmented in 3D using the following approach. First, we propagate the 2D binary mask of the CT along the Z-direction. This extended 3D mask is used to extract the 3D intensity voxels of the CT from the Z-stack images using a simple image masking operation. Next, Otsu thresholding was applied to the extracted 3D voxels followed by a morphological closing operation, to clean-up isolated voxels, for generating the 3D binary masks for CTs.

### 2.5. Statistical Analysis

All statistical measurements and graphics was generated using R version 3.3.2, 64-bit with ggplot2 graphics package (version 2.2.1, [27]). P-values were calculated using either the Wilcoxon-Mann-Whitney test [28] or the two sample Kolmogorov-Smirnov (KS) test [29, 30].

The number of CTs detected was calculated on a per chromosome basis in two biological replicates containing 3 technical replicate wells using R and plotted using Microsoft Excel. For nuclear volume and area as well as CT area and volume box plots, CT size were graphed on a per CT per karyotype basis in technical triplicates. For *Xist* pixel box plot, *Xist* was measured on a per X CT basis in triplicates. For centroid normalized radial positioning, distance from the border and nuclear spatial positioning were calculated using KNIME. Statistical analysis of *Xist* staining was done in technical triplicates.

### 2.6. Data and Software Availability

KNIME analysis workflows and R scripts are available on Github (https://github.com/CBIIT/Misteli-Lab-CCR-NCI)

## 3. Results

### 3.1. High-throughput Detection of CTs

We sought to develop an assay for the rapid, robust, and accurate identification of CTs at high-throughput to overcome the significant labor required for the acquisition, detection, and analysis of sufficiently large numbers of CTs for statistical analysis of their organization and localization (Fig. 1A, B). Our experimental pipeline includes staining of CTs in a 384-well plate format using chromosome paint FISH probes that were generated as previously described (see Materials and Methods). To adapt the standard CT FISH protocol incubation volumes, incubation times and washing steps were empirically optimized (see Materials and Methods). We applied this assay to the detection of human chromosomes 18 (HSA-18), X (HSA-X), and Y (HSA-Y) in normal human primary skin fibroblasts from female (XX) and male (XY) individuals (Fig. 1B). Typically, 12-13 fields per well were imaged per karyotype acquiring ~4000/cells per well using a high-throughput spinning disk confocal microscope. For image analysis, we used a custom image analysis workflow based on the software KNIME (see Materials and Methods) to identify nuclei with high precision using a seed watershed, ultrameric contour map for nucleus segmentation, followed by a random forest (RF) classifier for filtering out overlapping and mis-segmented nuclei. We typically analyzed a minimum of 1600 cells in two biological replicates containing 3 technical replicate wells per karyotype (Sup Fig. 1). A separate customized KNIME workflow was used to first detect CTs within the segmented nuclei using an undecimated wavelet-based spot/vesicle detection approach, and to then filter out background (non-CT) regions using a separate RF classifier. In addition, this workflow calculates various geometric and intensity-related features of the segmented CTs, including chromosome area and volume, as well as their position (only for 2D analysis) with respect to the nucleus, expressed either as a normalized Euclidean distance transform (nEDT) of the CT’s centroid or as the percent of the segmented CT region in each of five equidistant shells (Fig.1C).

Using this assay, we measured the expected number of HSA-X in a majority of cells from XY (90 +/- 6.0% SD) and XX (77 +/- 1.7% SD) cells in 2D (Fig. 1D). Similarly, HSA-Y was correctly detected in 88 +/- 2% SD of XY fibroblasts (Fig. 1D). Two copies of HSA-18 were detected in 70 +/- 1.7% SD of the cells in XY cells and 68 +/- 2% SD in XX cells. In many cases where the number of CTs detected was less than the expected (under-detection of CTs), it was not due to limitations in the segmentation algorithm since fewer copies of the corresponding chromosomes were also detected by visual inspection in those nuclei. In some cases, under-detection was due to clustering of the two homologue chromosomes and in others due to incomplete FISH hybridization (Sup Fig. 2). Less commonly, we observed that the number of CTs detected was greater than expected (over-detection of CTs), which was in part due to a combination of complex CT structure, the wrapping of CTs around nuclear structures (i.e. nucleoli), or the sub-optimal binding of chromosome paint probes to CTs, resulting in apparent fragmentation of the signals. For all further analysis, nuclei that contained an incorrect number of detected CTs for each channel were excluded from data analysis.

### 3.2. 2D vs 3D analysis of CT features

To determine how similar the detection of CTs in 2D compared to 3D, we systematically compared the percent of correctly segmented CTs using either 2D or 3D image analysis (Fig. 2A). For 2D analysis, 9-10 Z-planes were imaged, maximally projected in 2D and used to segment CTs using the undecimated wavelet-based spot detection method (see Materials and Methods). For 3D segmentation of CTs, we propagated the 2D binary mask along the Z-direction to extract 3D intensity voxels of the CT and then applied Otsu thresholding on these voxels to generate the 3D binary masks for the CTs (see Materials and Methods). We found remarkable correspondence in the number of detected CTs for HSA-18, X, and −Y using either 2D or 3D analysis (Fig. 2A). We used 2D and 3D analysis, respectively, to calculate nucleus area and volume of XX and XY cells (Fig. 2B). The 2D nucleus area of XY cells was slightly larger than that of XX cells (7107 vs 7369 average pixels per nucleus, 3.7% difference, Wilcoxon test p-value=3.5e-21) (Fig. 2B, top left), and similarly we detect a 3.7% increase in nuclear volume of XX nuclei compared to XY nuclei when analyzed in 3D (Wilcoxon test p-value =1.4e-11) (Fig. 2B, top right) in roughly 5,000 nuclei. With regards to CT features in male skin fibroblasts, as expected, the CT area and volume of HSA-X was the largest (2D: 225 +/- 23 pixels SD, 3D: 1982 +/- 183 pixels SD), HSA-Y the smallest (2D: 160 +/-3 pixels SD, 3D: 1380 +/- 51 pixels SD) and HSA-18 intermediate (2D: 199 +/- 9 pixels SD, 3D: 1660 +/- 64 pixels SD). The overall trends in our 2D measurements were replicated in our measurements in 3D of nuclear size and CT size (Fig. 2B and 2C), and show agreement between 2D and 3D analysis as shown previously [31]. Given the reduced computing time in 2D when compared to the similar analysis task in 3D, all subsequent image analysis of nuclei and CTs were done using 2D analysis.

### 3.3. Measurement of Xa and Xi chromosome size

We then used HiCTMap to probe the size variations of the HSA-X in populations of XY and XX fibroblasts. We observed that the HSA-X area in XY fibroblasts is significantly larger (median = 254 +/- 23 pixels) compared to XX fibroblasts (median = 212 +/-1 pixels SD) (Wilcoxon test p-value: 1.74e −11) as expected, due to inactivation and compaction of one of the X chromosomes in female cells (Fig. 3A) [32-35]. To distinguish between the two HSA-X in female cells, we ranked the CTs by size on a per nucleus basis (Fig.3B). The area of the larger X CT in XX fibroblasts was comparable to the single X CT in XY cells (256 +/- 23 SD vs. 254 +/- 3 pixels SD) (Wilcoxon test p-value > 0.05), whereas the smaller X CT which in most cases corresponds to the inactive X (see below), showed a 31% decrease in area compared to the larger X (176 +/- 2 pixels SD) (Wilcoxon test p-value: 2e-89) (Fig. 3B).

The well characterized mechanism of X chromosome inactivation (XCI) is initiated and maintained by the *Xist* lncRNA [36, 37]. *Xist* coats the inactive X chromosome (Xi) in cis and recruits repressive epigenetic markers, while the active X chromosome (Xa) is primarily used as the transcribed chromosome [38, 39]. To visualize and unambiguously identify the inactive X chromosome, XX cells were processed for sequential DNA/RNA FISH using chromosome-paint probes to label the X CT and fluorescently labeled oligonucleotides to target *Xist* RNA (see Methods). This approach enables high-throughput visualization of simultaneously labelled X chromosomes and *Xist* RNA in the same nucleus (Fig. 3C). Any nucleus containing more than one *Xist* focus was excluded from further analysis. Figure 3D shows that the Xa contains no *Xist* staining compared to the Xi chromosome, and the Xi is typically smaller compared to Xa (Fig. 3D, E). As expected, while we find a robust single *Xist* RNA signal in nuclei from XX cells, no *Xist* RNA signals are detected in XY cells (data not shown). In line with our size analysis, we find that CTs that lack *Xist* label are typically larger (225 pixels +/- 11 SD) compared to *Xist* containing CTs (164 +/- 14 pixels SD) (Wilcoxon test p-value: 8.23e-121, Fig. 3E). These observations demonstrate accurate detection and the ability to distinguish Xi and Xa based on size and *Xist* staining and they establish the feasibility of using HiCTMap to combine DNA and RNA FISH in high-throughput imaging approaches.

### 3.4. Chromosome Positioning Analysis

We finally applied HiCTMap to determine the nuclear positioning of HSA-18, -X, and -Y in XY fibroblasts. We used two approaches to determine chromosome positioning. First, we determined the centroid of the segmented CT defined as the geometric center of the segmented CT and measured its normalized radial distance from the nucleus border using Euclidean distance transformation as previously described [40]. The distribution of the centroid radial distance for each chromosome was distinct (Fig. 4A). HSA-X showed a strong preferential distribution to the periphery of the nucleus in XY cells (median normalized radial distance = 0.34), whereas HSA-Y exhibited a slight preference for positioning to the center of the nucleus (median normalized radial distance = 0.58) and HSA-18 showed an even distribution of positions (0.48) (Fig.4A). Comparison of the radial positioning of the centroid of Xa and Xi in XX fibroblasts, showed a small, but significant, more peripheral position of the *Xist*-marked Xi centroid (median normalized radial distance = 0.25) compared to the Xa (median normalized radial distance = 0.29) (KS test p-value: 2.24e-07) (Fig.4B). Measurements of center of gravity, which accounts for intensity values in the CT, gave similar results (data not shown).

Since using centroids as a proxy for the spatial localization of an entire CT could potentially generate misleading results, given the irregular and diffuse structure of chromosome territories, we also used an alternative method to more faithfully determine chromosome location. In this approach, each nucleus is divided into five equidistant shells, and the percentage of the CT falling in each of the equidistant shells is measured [32, 41]. When chromosome positioning was measured using equidistant shells, we observed similar results to the centroid analysis: a strong preferential positioning of the HSA-X to the periphery of the nucleus with the highest average percent in shells 1 and 2 (30% and 32%, respectively), a slight preference for positioning of HSA-Y to the center of the nucleus with the highest average percent in shells 3 and 4 (24% and 27%, respectively) and even positioning of HSA-18 with the highest average percent in shells 2 and 3 (24% and 25%, respectively) (Fig.4C). Similar to the results obtained with centroids, in XX fibroblasts, both Xa and Xi were peripherally positioned, and Xa was slightly more internal (38% in shell 5) compared to Xi (42% in shell 5) (Fig.4D).

Taken together, these results establish a high-throughput imaging pipeline for the detection of CTs and the determination of structural features of chromosomes including area, size and radial position. In addition, we show that chromosome painting in a high throughput format can be used to detect multiple chromosome combinations in the same experiment and can be combined with RNA FISH. We suggest that HiCTMap is a versatile tool for the study of chromosomes at a high-throughput scale.

## 4. Discussion

Here, we describe a systematic and quantitative method for the detection of multiple chromosome territories using high-throughput imaging. Our approach, referred to as HiCTMap, enables collection of imaging data on large numbers of cells and quantitative measurements of chromosome features including size and positioning using automated image analysis tools. HiCTMap overcomes the constraints of standard CT analysis, particularly the reliance on relatively small numbers of cells due to technical limitations in sample preparation and extensive imaging time using conventional microscopy approaches.

Due to its high-throughput nature, HiCTMap is well suited to acquire images for several thousands of cells per experimental condition in a short time, typically about seven minutes per well. In contrast to most FISH approaches that rely on visual inspection of relatively small sample numbers and require significant user input, automated imaging of thousands of cells per sample generates a more faithful representation of the frequency of chromosomal architecture and positioning in the population and eliminates user biases in selection of nuclei for quantitative analysis. We demonstrate that chromosome painting in high-throughput can robustly detect the expected number of CTs simultaneously in three channels. We also establish the accuracy of HiCTMap in measuring the position of chromosomes using two methods of radial distance measurements, equidistant shells and centroid and we show its compatibility with RNA FISH.

As previously reported [31] for traditional imaging methods, we find little differences when analyzing CTs in 2D or 3D. The number of chromosomes detected in male (XY) and female (XX) cell populations was similar in 2D and 3D analysis as was nuclear size and chromosome size. The relatively large size of chromosomes in nuclear space provides a lower threshold for accurate detection, segmentation, and analysis than for the precise and meticulous accuracy of single gene FISH in 2D versus 3D [31]. It is worth noting that use of higher-magnification imaging provides better resolution of chromosome architecture, however, 60X image acquisition yielded similar results to our standard 40X magnification used and provided no further benefits to measurements in 2D or 3D (data not shown).

HiCTMap is sufficiently sensitive to detect expected differences in X chromosome size in XY and XX cells, due to the presence of a mixed population of smaller Xi and larger Xa chromosomes. While the population of X chromosomes in XY cells is relatively homogeneous, a more heterogeneous distribution of X chromosome size was found in XX cells, consistent with the well-established smaller size of the inactive X chromosome [32]. The identity of the consistently smaller X chromosome as the Xi was confirmed by sequential *Xist* RNA FISH and chromosome painting. Chromosome painting in high-throughput in combination with *Xist* RNA FISH offers advantages over conventional RNA FISH followed by imaging and subsequent DNA FISH, since it allows for immediate acquisition of both RNA and DNA FISH in one imaging step, eliminating the possibility of misaligning or incorrectly identifying nuclei when plates or cover slips are removed for DNA FISH and re-imaged. In addition, in HiCTMap, multiple chromosomes can be observed in combination with the *Xist* RNA probe, thus making the method suitable for use in basic research applications to probe aneuploidies of the genome.

Furthermore, we show robust detection by HiCTMap of chromosome positioning using both centroid and equidistant shell radial positioning approaches. We also detected differences in nuclear positioning of Xi and Xa using combined *Xist* RNA and CT FISH. Similar trends were observed by measuring the centroid location or equidistant shell radial positioning. Our findings are in line with the reported observation of a more peripherally positioned Xi and its proposed anchored to the nuclear lamina [42]. Unlike previous observations where a much greater difference in positioning is observed between the two X chromosomes [17, 19], our results may be different due to a different cell line used or the large number of nuclei measured that represents the population more faithfully.

The limitations of HiCTMap are relatively minor. First, CT segmentation requires user input to determine the ground truth necessary to implement the random forest filtration. This step typically requires 30 minutes of training and harbors the possibility of biases, which, however, can be eliminated by careful supervision of the user and determination of the ground truth by multiple users. Second, our imaging conditions were not optimized for maximal resolution in 3D. However, this seems unlikely as we observe similar segmentations results for all measured chromosomes after increasing the magnification from 40X to 60X as previously stated (data not shown).

Taken together, we describe here HiCTMap, a method for chromosome painting in high-throughput. Given its versatility and its compatibility with RNA FISH, we anticipate that HiCTMap will be of considerable use in future analysis of chromosome organization and function.

## Acknowledgement

The authors thank the CBIIT Server Team, NCI, National Institutes of Health (NIH) and the High Performance Computing Group, CIT, NIH for computational support. This research was supported by funding from the Intramural Research Program of the National Institutes of Health (NIH), National Cancer Institute, and Center for Cancer Research.

## FIGURE LEGENDS

**Figure 1. HiCTMap Imaging, Analysis, and Measurement**.

A) HiCTMap pipeline. Cells are cultured in 384-well imaging plates and DNA FISH is carried out using chromosome paints, followed by automated image acquisition using high-throughput microscopy. Image analysis by KNIME segments the nuclei and detects CTs in three channels. CT features are measured and plotted using the R software. B) CT signal detection. Representative maximal projections of images acquired in three channels. Maximal projections of 40X confocal images z-stacks in 4 channels of XX and XY skin fibroblast stained with DAPI (Blue- 408) and chromosome paint probes (X-Green (Alexa488), 18- Red (Dy505), and Y- FarRed (Dy651)). Scale bar: 20 μm. C) KNIME automated image segmentation pipeline. Nuclei are segmented using the DAPI (4’,6-diamidino-2-phenylindole) channel, thus generates a segmentation mask. The fluorescent channel of each chromosome paint probe is then used to segment CTs of HSA-18, -X, and -Y. Positioning of segmented chromosomes using normalized distance transformation and equidistant shells is calculated using the KNIME image analysis pipeline (see Materials and Methods for details). D) Histogram showing the number of CTs detected in XX and XY nuclei using 2D segmentation. Values represent averages from two biological replicates containing 3 technical replicates ± SD.

**Figure 2. 2D vs. 3D image analysis of segmented CTs and nuclear size**

A) Histograms showing the number of chromosomes 18, X, and Y per cell in XY nuclei using 2D or 3D analysis. Values represent averages from three experiments ± SD. B) Top left: Box plots of measured nuclear area of XX and XY cells using 2D nuclear segmentation. Top right: Box plots of measured nuclear volume of XX and XY cells using 3D nuclear segmentation. C) Bottom left: Box plots of measured CT area of chromosomes 18, X, and Y in XY cells using 2D CT segmentation. Bottom right: Box plot of measured CT volume of chromosomes 18, X, and Y in XY cells using 3D CT segmentation method. Boxes show the 25th, 50th (median) and 75th percentile of the distributions and whiskers extend to 1.5 x inter-quantile range (IQR), outliers are represented as dots. Notches indicate the estimated 95% confidence interval of the median (CI). Distributions represent data from approximately 5,000 nuclei.

**Figure 3. Detection and measurement of Xa and Xi**

A) Box plots of X CT area per nucleus in XX and XY cells. B) Box plots of X CT area of the largest (1) and second (2) largest ranked X CT per nucleus in XX and XY cells. C) Representative images of combined DNA-RNA FISH (Scale bar: 20 μm). D) Scatter plot of X CT area in pixels vs. the segmented *Xist* signals normalized to nuclear size in XX cells (Xa= active X chromosome, Xi= inactive X chromosome). E) Box plots of CT area in Xi vs. Xa per nucleus in XX cells on a per CT basis. Boxes show the 25th, 50th (median) and 75th percentile of the distributions and whiskers extend up to 1.5* inter-quantile range, outliers are shown as dots. Notches indicate the estimated 95% confidence interval of the median (CI). Distributions represent data from approximately 2,500 nuclei.

**Figure 4. Radial positioning of CTs**

A) Density curves for normalized radial distance distributions for the indicated chromosomes. B) Density curves for normalized radial distance distributions of Xa and Xi. C) The radial 2D distribution of HSA-18, -X, and -Y coverage in 5 concentric equidistant nuclear shells. D) The radial distribution of Xi and Xa coverage in 5 concentric equidistant nuclear shells. Values represent averages from three technical replicates ± SD. Distributions represent data from at least 2,500 CTs.

**Supplementary Figure 1.**

Histogram of the number of detected nuclei per well in XX and XY fibroblast using HiCTMap. Values represent averages from 3 technical replicates ± SD.

**Supplementary Figure 2.**

Representative DAPI, Red fluorescent channel (Excitation: 561 nm) of chromosome paint probe, and merge images of two HSA-18 chromosomes co-localizing.

